# Genomic consequences of isolation and inbreeding in an island dingo population

**DOI:** 10.1101/2023.09.15.557950

**Authors:** Ana V. Leon-Apodaca, Manoharan Kumar, Andres del Castillo, Gabriel C. Conroy, Robert W Lamont, Steven Ogbourne, Kylie M. Cairns, Liz Borburgh, Linda Behrendorff, Sankar Subramanian, Zachary A. Szpiech

**Affiliations:** Department of Biology, Pennsylvania State University, PA, USA; School of Science, Technology & Engineering, University of the Sunshine Coast, 1 Moreton Parade, Petrie, Queensland, Australia; School of Science, Technology & Engineering, University of the Sunshine Coast, 90 Sippy Downs Drive, Sippy Downs, Queensland, Australia; Centre for Bioinnovation, University of the Sunshine Coast, 90 Sippy Downs Drive, Sippy Downs, Queensland, Australia; Evolution & Ecology Research Centre, School of Biological, Earth and Environmental Sciences, UNSW Australia, Sydney NSW 2052, Australia; Centre for Ecosystem Science, School of Biological, Earth and Environmental Sciences, UNSW Australia, Sydney NSW 2052, Australia; Queensland Parks and Wildlife Service, Department of Environment & Science, K’gari, Australia; Institute for Computational and Data Sciences, Pennsylvania State University, PA, USA

## Abstract

Dingoes come from an ancient canid lineage that originated in East Asia around 8000-11,000 years BP. As Australia’s largest terrestrial predator, dingoes play an important ecological role. A small, protected population exists on a world heritage listed offshore island, K’gari (formerly Fraser Island). Concern regarding the persistence of dingoes on K’gari has risen due to their low genetic diversity and elevated inbreeding levels. However, whole-genome sequencing data is lacking from this population. Here, we include five new whole-genome sequences of K’gari dingoes. We analyze a total of 18 whole genome sequences of dingoes sampled from mainland Australia and K’gari to assess the genomic consequences of their demographic histories. Long (>1 Mb) runs of homozygosity (ROH) — indicators of inbreeding — are elevated in all sampled dingoes. However, K’gari dingoes showed significantly higher levels of very long ROH (>5 Mb), providing genomic evidence for small population size, isolation, inbreeding, and a strong founder effect. Our results suggest that, despite current levels of inbreeding, the K’gari population is purging strongly deleterious mutations, which, in the absence of further reductions in population size, may facilitate the persistence of small populations despite low genetic diversity and isolation. However, there may be little to no purging of mildly deleterious alleles, which may have important long-term consequences, and should be considered by conservation and management programs.

**SIGNIFICANCE:** A long-standing question in conservation genetics is whether long-term isolation and elevated levels of inbreeding always leads to inevitable population extinction. Here we conduct the first-ever whole-genome analysis of a population of dingoes living in long-term isolation on an island off the coast of Australia (K’gari). We show that these animals are beset by very low genetic diversity, likely the result of extensive inbreeding, and an elevated number of deleterious homozygotes. However, our results suggest that these dingoes are likely purging highly deleterious alleles, which may have allowed them to persist long term despite their extremely small population (<200 individuals).

## INTRODUCTION

Dingoes have a complex history in Australia, as they have been simultaneously declared a native species and an agricultural pest dependent on jurisdiction and context. While they are protected in parts of the Australian Capital Territory, Victoria, Queensland and the Northern Territory as a native species, ‘wild dogs’, including dingoes, have been declared a pest or restricted matter in all Australian states and territories for posing a risk to livestock (Brink et al. 2019).

The dingo plays an important ecological role in maintaining the ecosystems of Australia, as they have been the apex terrestrial predator since the extinction of the thylacine (Glen et al. 2007; Johnson et al. 2007; Letnic et al. 2009; Wallach et al. 2010; Letnic et al. 2011; Letnic et al. 2012). Research using mtDNA and Y-chromosome data has identified a close relationship between dingoes, New Guinea singing dogs (NGSD), and Asian village dogs, suggesting that the ancestors of dingoes originated in East Asia (Savolainen et al. 2004; Oskarsson et al. 2012; Ardalan et al. 2012; Sacks et al. 2013; Cairns and Wilton 2016; Cairns et al. 2017). Archeological evidence and genomic data estimate the divergence time between wolves and early dogs to be ∼15,000-36,000 years BP (Germonpré et al. 2009; Germonpré et al. 2012; Larson et al. 2012; Skoglund et al. 2015; Frantz et al. 2016; Freedman and Wayne 2017), and ∼5,000-11,000 years BP between early dogs and dingoes (Savolainen et al. 2004; Oskarsson et al. 2012; Sacks et al. 2013; Cairns and Wilton 2016; Bergström et al. 2020; Surbakti et al. 2020; Cairns 2021; Field et al. 2022). Although dingoes are descended from a dog-like ancestor, they form a distinct and separate lineage from domestic dogs, wolves and other canids (Crowther et al. 2014; Smith et al. 2019; Cairns 2021; Field et al. 2022).

At least two distinct dingo lineages have been identified in the mainland, one in the northwest and one in the southeast, alongside one offshore lineage on K’gari (formerly known as Fraser Island) (Cairns and Wilton 2016; Cairns et al. 2017; Cairns et al. 2018; Greig et al. 2018; Zhang et al. 2020). K’gari is located off the coast of Queensland, Australia, and has been listed as a UNESCO World Heritage site since 1992. In addition to being the world’s largest sand island, it is also home to a small, protected (Nature Conservation Act 1992) population of dingoes of ∼70-170 individuals that are genetically distinct from mainland dingoes (Brown et al. 2011; Conroy et al. 2021). In contemporary times dingoes are known for being an infamous tourist attraction for the over 400,000 tourists that visit K’gari each year (Wardell-Johnson et al. 2015), which has led to human-wildlife conflict (O’Neill et al. 2017; Behrendorff 2021). In response to these incidents, management strategies have been implemented throughout the island, including non-lethal options as well as the removal of dingoes via lethal control, although the latter is currently a last resort action and only occurs around high-use visitor areas (Behrendorff 2021). K’gari dingoes are of particular interest to conservationists, as well as various stakeholders in Australia, as there are concerns regarding their persistence due to low genetic diversity and inbreeding given it is a small, isolated population (Cairns et al. 2018; Conroy et al. 2021).

Isolated populations often experience reduced fitness resulting from increased levels of homozygosity and a high load of deleterious mutations across their genomes. There have been several instances in which these factors have led to inbreeding depression and a high risk of extinction. Such is the case of the iconic Isle Royale wolves, which are highly inbred gray wolves located on an island in Lake Superior, in northern United States (Hedrick et al. 2019; Robinson et al. 2019). Other examples of isolated populations in the wild that have experienced inbreeding depression are Florida Panthers (Roelke et al. 1993) and Scandinavian gray wolves (Liberg et al. 2005; Räikkönen et al. 2006; Kardos et al. 2018). In other instances, small, isolated populations have managed to persist with low genetic diversity for generations, without experiencing inbreeding depression. Such is the case of Channel Island foxes (Robinson et al. 2016; Robinson et al. 2018), Appenine brown bears (Benazzo et al. 2017), African mountain gorillas (Xue et al. 2015), Alpine ibex (Grossen et al. 2020), and Isle Royale moose (Kyriazis et al. 2023). It has been suggested that the purging of strongly deleterious, recessive mutations may facilitate the persistence of long-term small populations (Xue et al. 2015; Robinson et al. 2018; Grossen et al. 2020; Wootton et al. 2023).

Only a handful of studies have analyzed the genetics of K’gari dingoes. While these studies have found low genetic diversity and high levels of inbreeding on K’gari dingoes (Cairns and Wilton 2016; Cairns et al. 2018; Conroy et al. 2021), more research is needed to understand the consequences of inbreeding and isolation on this population. Moreover, there is a need for new and well-informed conservation strategies for dingoes in Australia, given that current strategies do not take into account these genetic factors. To our knowledge, there is only one whole-genome sequence from a K’gari dingo individual that is publicly available (Zhang et al. 2020). In this work we include five newly sequenced samples, and we look further into the genetics of K’gari dingoes to understand the consequences of their demographic history on the distribution of deleterious variation, and the implications of small population sizes and isolation, to help inform conservation efforts to ensure their long-term persistence.

## RESULTS

### Canid Sampling, Relatedness and Population Structure

Previous analysis on dingoes suggest there are at least three distinct lineages of dingoes in Australia, with one being located on the northwest of the mainland, another in the southeast of the mainland, and the third one offshore on K’gari (Fraser Island) (Cairns and Wilton 2016; Cairns et al. 2017; Cairns et al. 2018). To expand our understanding of genetic diversity patterns in the K’gari population, we generated high-quality, whole-genome sequences of five dingoes with average read depth between 12.01X and 12.56X sampled from the island between 2011 and 2013. Although there is a dingo reference genome (ASM325472v1), the newly sequenced samples in this study were aligned to the dog reference genome (CanFam3) to take advantage of its fully assembled chromosomes and better gene annotations. We include previously published whole-genome sequences from dingoes from northwest and southeast mainland Australia, from wolves, village dogs, and New Guinea singing dogs (NGSD), which are a close relative of dingoes. All canid reads in this study were aligned to the dog reference genome (CanFam3).

We identified close relatives in K’gari dingoes in two ways. First, we looked at the proportion of sites at which pairs of individuals share 0, 1 or 2 alleles that are identical-by-state (IBS). We expect Parent/Offspring to share at least one allele at a locus identical-by-descent, thus having low values of IBS0. We expect full-siblings to share 2 alleles identical-by-descent for approximately 25% of loci, and we expect second-degree relatives and unrelated individuals to have higher estimates of IBS0. We also used the KING algorithm (Manichaikul et al. 2010) to infer kinship coefficients for all dingo individuals. We identified three pairs of Parent/Offspring relatives, four pairs of full-siblings relatives, a pair of avuncular relatives, and seven pairs that include third-degree relatives and unrelated individuals. Furthermore, we found that none of the K’gari dingo individuals are closely related to dingo individuals from the mainland, and that the mainland dingoes in this study are not closely related to one another (supplementary fig. S1; supplementary table S1).

We computed pairwise allele sharing dissimilarities to assess population structure and generated multidimensional scaling (MDS) plots from the dissimilarity matrix. In MDS, individuals with small allele sharing dissimilarities tend to group together. In fig. 1A we can observe distinct taxonomic groupings, with the close relationship between the dingo and NGSD lineages clearly captured in this analysis. Individuals within the wolf and village dog populations were sampled from different regions around the world (see supplementary table S2), hence the relatively loose clustering among these individuals. Interestingly, we find that the village dog from Vietnam seems to be genetically more similar to mainland dingoes than the village dog from Papua New Guinea (fig. 1A). While Papua New Guinea is geographically closer to Australia than Vietnam, similar observations regarding shared ancestry between Vietnamese village dogs and dingoes have been reported by Cairns et al. (Cairns et al. 2023). A possible explanation regarding the observed genotype differences among these canids may be explained by Vietnamese village dogs having retained the ancestral alleles that also exist in the dingo lineage while village dogs from Papua New Guinea have not, possibly because of their different demographic histories or genetic swamping from contemporary domestic dogs.

**Fig. 1.**
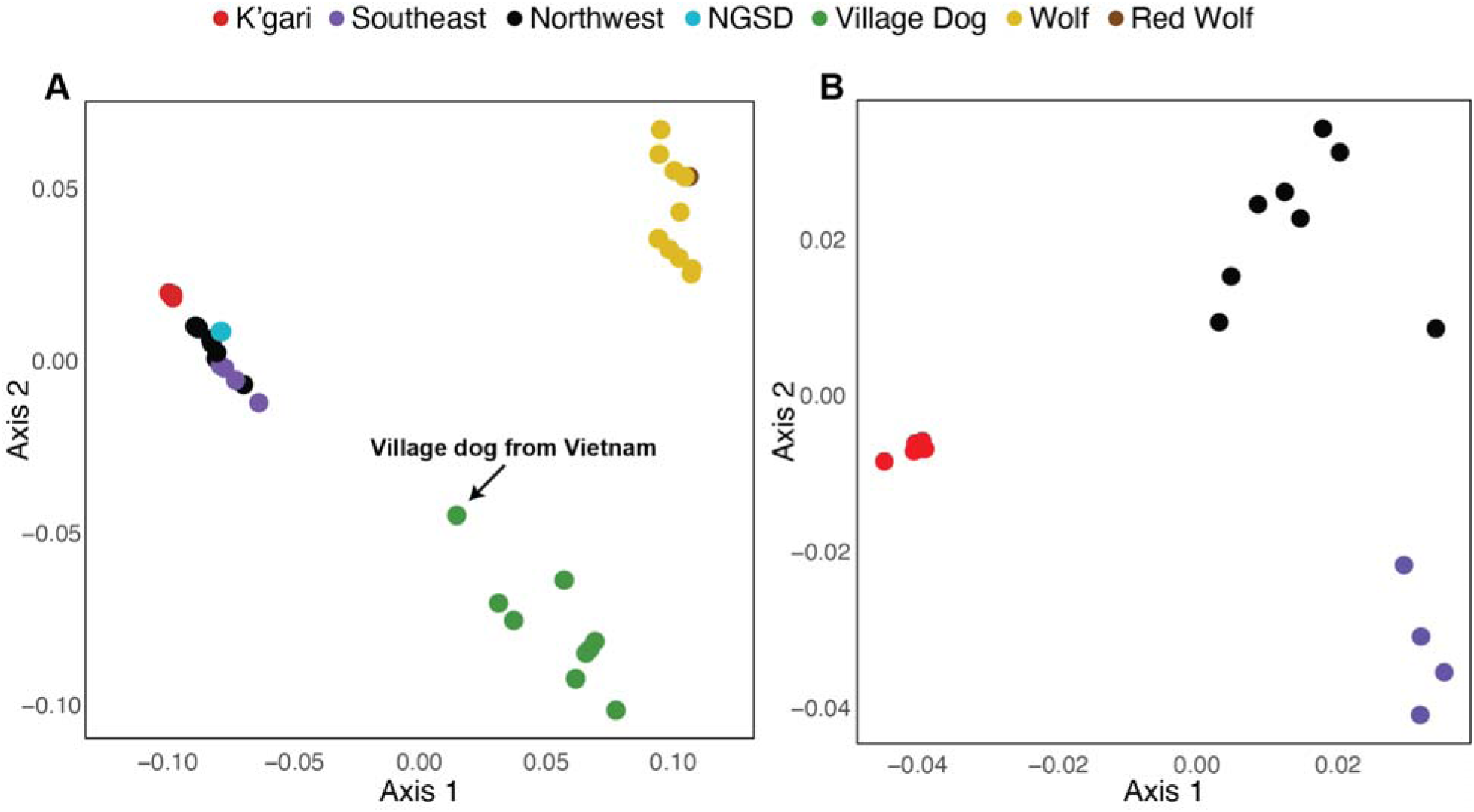
Reduced dimensional representation of sample clustering based on multidimensional scaling of the allele sharing dissimilarity matrix. First two major axes of variation plotted for (A) all 41 canids included in this study and (B) 18 dingo individuals.

Although dingoes from the southeast and northwest overlap with each other (i.e., they are genetically more similar), we observe K’gari dingoes form their own separate subgroup (fig. 1A). To look more into detail at the allele sharing differences between the three dingo populations, we recomputed the MDS on the matrix of allele sharing dissimilarities containing only these samples. In fig. 1B we can observe geographic based clustering, with K’gari dingoes maintaining a tighter and more distinct group compared to dingoes in the southeast and northwest.

### Genetic diversity between and within canid populations

To characterize genetic diversity patterns within and between canid populations we analyzed the distribution of alleles across populations. We used ADZE (Szpiech et al. 2008) to compute estimates of allelic richness, private allelic richness, and the number of alleles shared by pairs of populations (i.e., alleles that are found only in those populations and nowhere else). We observed reduced levels of genetic diversity in mainland and K’gari dingoes, compared to large and outbred populations such as wolves and village dogs, with K’gari dingoes having the lowest genetic diversity of all the canid groups analyzed in this study (fig. 2A; fig. 2B). Furthermore, we found mainland dingoes share more alleles with village dogs than K’gari dingoes and village dogs (fig. 2C). Although hybridization with domestic dogs was previously highlighted as a main concern for dingo conservation (Newsome and Corbett 1985; Jones 1990; Corbett 2001; Wilton 2001; Elledge et al. 2008; Stephens et al. 2015), more recent studies have shown hybridization is uncommon (Cairns et al. 2020; Cairns 2021; Cairns et al. 2021; Cairns et al. 2023). This suggests that the alleles shared between dingoes and their common ancestors from Southeast Asia were likely lost in the bottleneck that founded the K’gari dingo population.

**Fig. 2.**
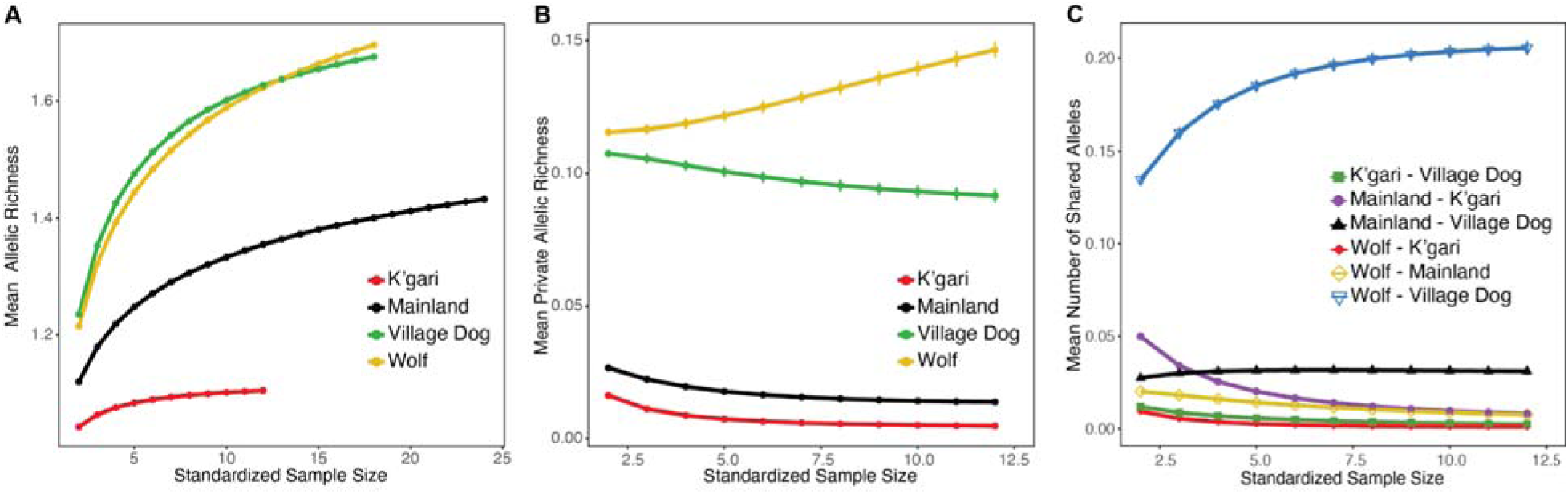
Genetic diversity estimates within and between canid populations. Error bars representing 95% confidence intervals for each line are too narrow to be visible. (A) Mean number of alleles per locus (allelic richness). (B) Mean number of private alleles per locus (private allelic richness). (C) Mean number of shared alleles per locus between pairs of populations.

### Inbreeding and homozygosity

To characterize the levels of inbreeding we looked at long contiguous stretches of homozygous regions in individual genomes that are shared identical-by-descent (IBD), also known as runs of homozygosity (ROH). We used GARLIC (Szpiech et al. 2017) to call ROH, which implements a likelihood-based approach to infer ROH. The length of ROH can inform on how recent the parental relatedness among individuals is, with long ROH more commonly arising in offspring descended from parents with a recent common ancestor. On the other hand, short ROH suggests a more distant common ancestor, as recombination has had more time to break apart the underlying IBD haplotypes (Kirin et al. 2010; Pemberton et al. 2012). The wolf and village dog individuals included in this study were sampled from large and outbred populations from different regions around the world (supplementary table S2), resulting in variation in the levels of homozygosity found in their individual genomes. Overall, we observed relatively low levels of homozygosity in village dogs and wolves, with ∼7% and ∼16% of their genomes being covered by ROH >0.5 Mbp, respectively (fig. 3). In contrast, we observed high levels of homozygosity in northwest and southeast mainland dingo populations, with ∼30% of their genomes being covered by ROH >0.5 Mbp. In K’gari dingoes we observed even higher levels of homozygosity, with ∼65% of their genomes being covered by ROH >0.5 Mbp, and ∼28% by ROH >5 Mbp (fig. 3), with the longest individual homozygous segment being 64.2 Mbp long (supplementary fig. S3).

**Fig. 3.**
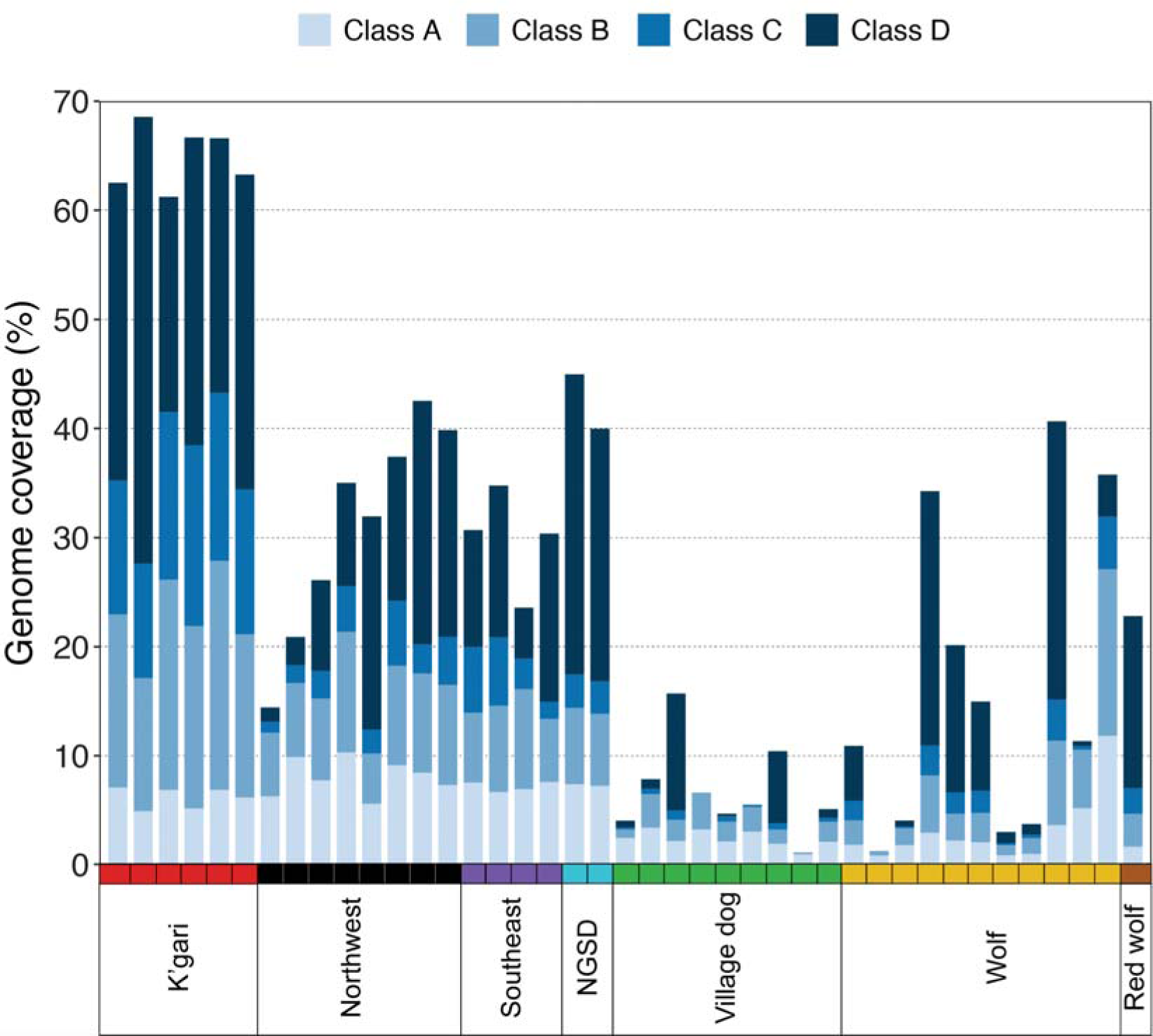
ROH coverage across individual genomes. The percentage of the genome covered by class A ROH (0.5-1 Mbp), class B ROH (1-3 Mbp), class C ROH (3-5 Mbp) and class D ROH (>5 Mbp) is shown on the Y-axis for each of the 41 canids included in this study.

We estimated inbreeding coefficients as given by ROH for each population (F_ROH_), which represents the proportion of the autosomal genome covered by ROH >1Mbp. Mean F_ROH_ of 0.59, 0.23, 0.23, were estimated for K’gari, Northwest, and Southeast dingoes, respectively, and 0.35 for NGSD. For village dogs and wolves we estimated mean F_ROH_ of 0.04 and 0.13, respectively. These results provide clear evidence for inbreeding, isolation, small population size, and a strong founder effect in K’gari dingoes.

### Patterns of deleterious variation and genetic load

To understand the effect of inbreeding and low genetic diversity on the distribution of deleterious variation, we looked at the burden of deleterious alleles in K’gari dingoes compared to mainland dingoes and other canids. Alternate alleles were predicted to be deleterious or neutral by taking into account sequence conservation and the effect of an amino acid substitution on protein function using Ensembl VEP (McLaren et al. 2010) and SIFT (Vaser et al. 2016) (see Methods). We found dingoes and NGSD carry a higher number of deleterious alternate-allele homozygotes compared to village dogs and wolves, although K’gari dingoes carry the highest number of deleterious alternate-allele homozygotes among all the canids included in this study (fig. 4A). Conversely, we observed K’gari dingoes carry the lowest number of deleterious and LoF alternate-allele heterozygotes compared to the other canids (fig. 4B; fig. 4D). Furthermore, we observed a significant difference in the number of alternate-allele LoF homozygotes, with K’gari dingoes carrying a higher number than northwest and southeast mainland dingoes (fig. 4C).

**Fig. 4.**
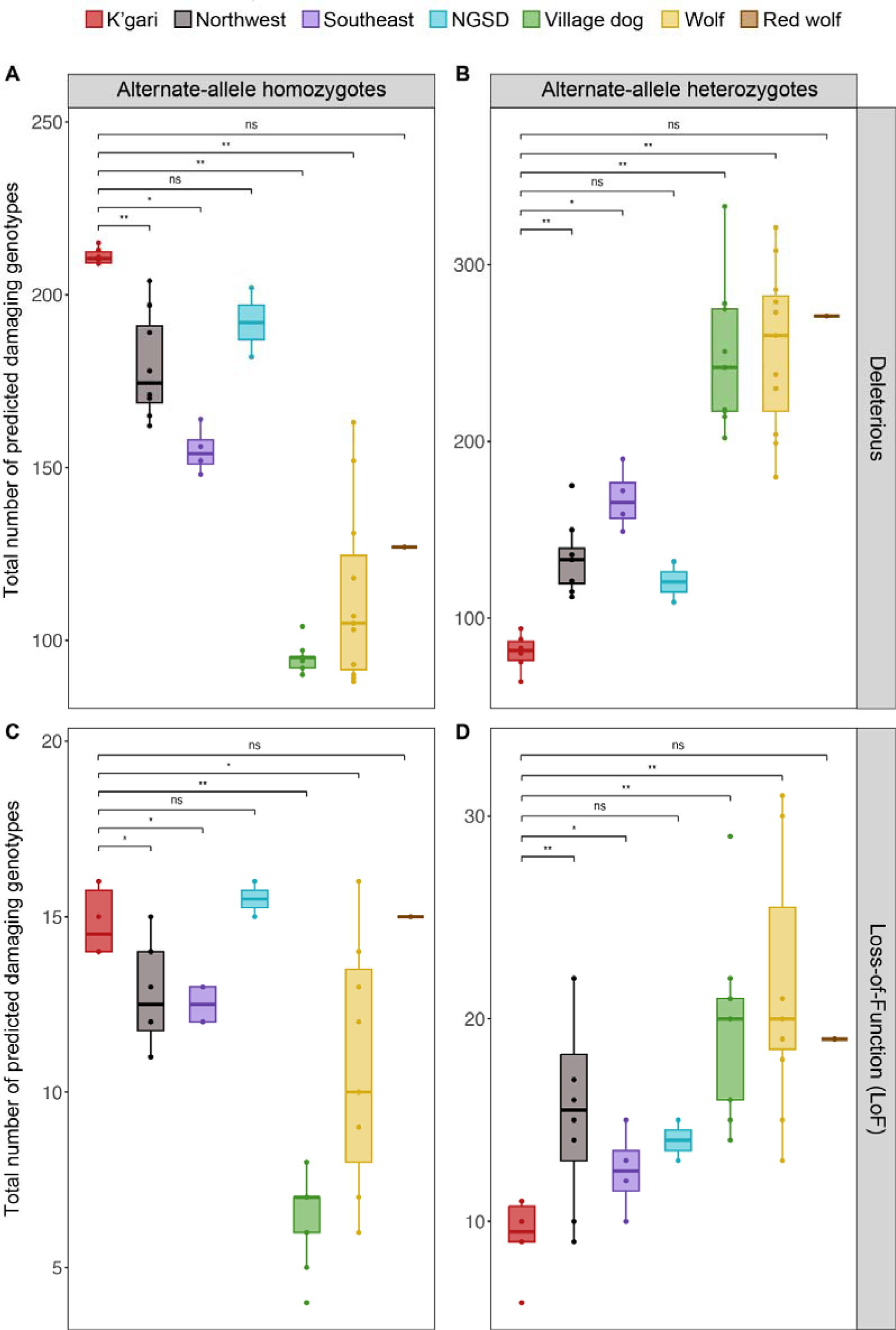
Total number of predicted damaging genotypes in each canid population. (A) Alternate-allele deleterious homozygotes. (B) Alternate-allele deleterious heterozygotes. (C) Alternate-allele LoF homozygotes. (D) Alternate-allele LoF heterozygotes. P-value significance by Wilcoxon test with K’gari dingoes as reference population. Significance levels of p-values: (*) p-value < 0.05, (**) p-value < 0.005, (ns) p-value > 0.05.

To refine our analysis of the differences in the burden of deleterious alleles between K’gari and mainland dingoes, we subset our deleterious and LoF alternate-allele calls to include only those alternate alleles which were also inferred to be the derived allele in dingoes. We observed patterns of deleterious derived homozygotes and heterozygotes stayed qualitatively similar to the non-polarized patterns (supplementary fig. S4A; (supplementary fig. S4B). For the LoF category, we did not find a significant difference in the number of derived homozygotes in K’gari and mainland dingoes (supplementary fig. S4C), however, the mean number of LoF derived heterozygotes in K’gari dingoes is significantly lower than in mainland dingoes (supplementary fig. S4D). Similar patterns have been observed in other small populations with substantial inbreeding, suggesting purging of LoF alleles, which are more likely to be strongly recessive variants than small-effect deleterious variants, may facilitate the persistence of small and isolated populations (Robinson et al. 2018; Grossen et al. 2020).

To test whether K’gari dingoes showed evidence of purging of deleterious variants we used the Rxy statistic, as implemented by Xue et al. (Xue et al. 2015), which compares the frequency of derived alleles of a particular category (e.g., deleterious) normalized by the frequencies of a set of putatively neutral variants, which is then compared across two populations, X and Y. When Rxy < 1 for a given site category, we consider this evidence of purging of those alleles in population X relative to Y, when Rxy > 1 we consider this evidence of enrichment of those alleles in population X relative to Y. When Rxy = 1, we consider this evidence of no difference of those alleles between X and Y. As our K’gari samples are closely related, we implemented a modified version of the Rxy statistic, which accounts for relatedness among K’gari dingoes (see Methods). For this analysis we looked at several allele categories: deleterious, tolerated, LoF, synonymous and nonsynonymous sites, and a random set of intergenic sites as a control. All Rxy calculations used an additional, independent, set of intergenic sites for normalization. We estimated the excess number of deleterious variants in K’gari dingoes (population X) with respect to mainland dingoes (population Y) and found evidence of purging of LoF alleles (i.e., putative strongly deleterious alleles), but little to no evidence of purging of mild/slightly deleterious alleles in K’gari dingoes (fig. 5). Although we observe a small enrichment in the rest of the categories, we hypothesize these results may be due to the small sample size and low genetic diversity in K’gari dingoes compared to mainland dingoes. To assess the effect of sample sizes on our results, we conducted this analysis using the same number of samples for each dingo population and found that results stayed very similar (supplementary fig. S5).

**Fig. 5.**
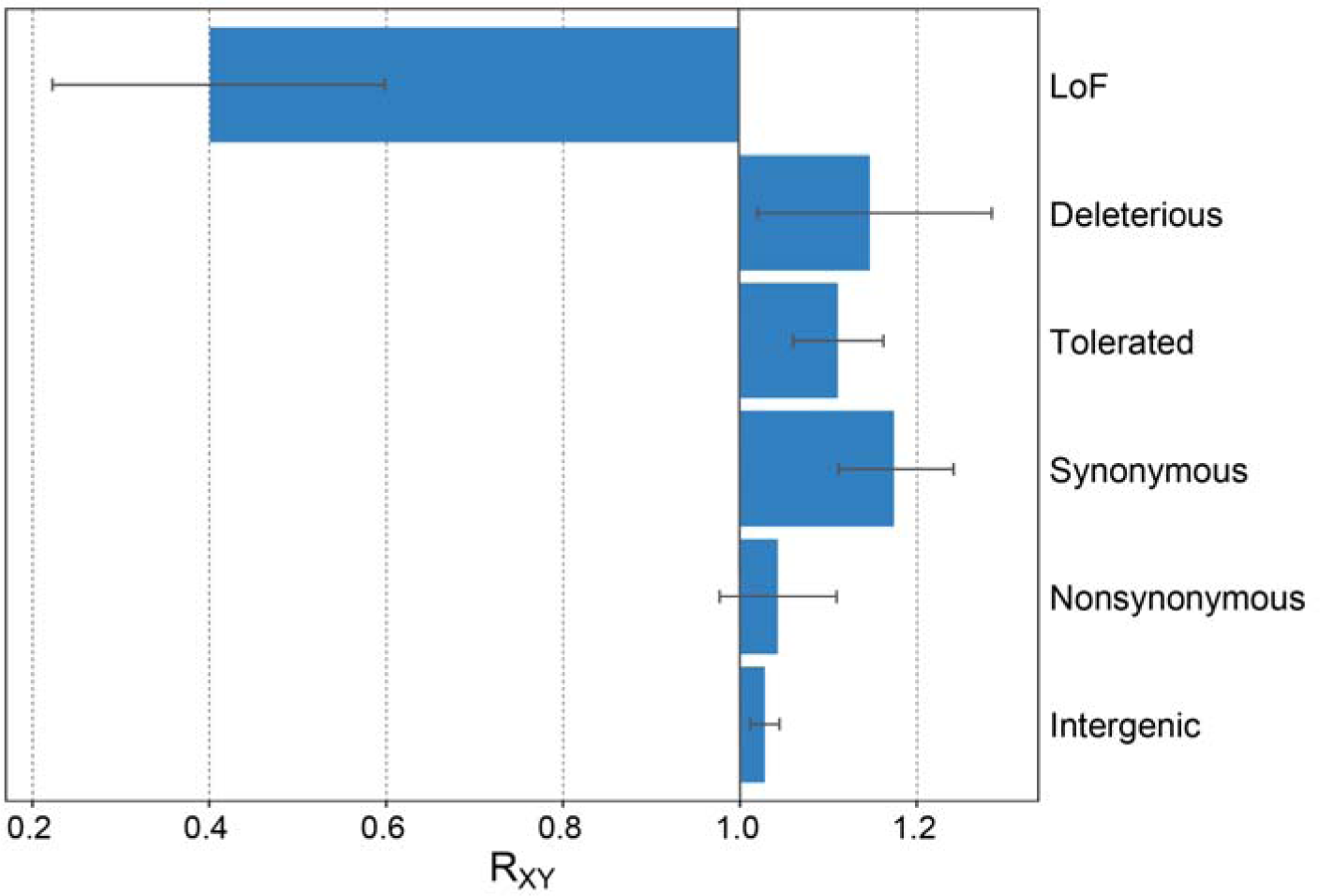
Kinship-weighted Rxy estimates for K’gari and mainland dingoes. Estimates represent the relative excess number of derived alleles for each category in K’gari dingoes (population X) relative to mainland dingoes (population Y). Error bars represent the 95% confidence interval from jackknife resampling.

## DISCUSSION

Inbred populations frequently experience inbreeding depression and subsequently face a high risk of extinction. However, some populations are able to persist in small numbers and with low genetic diversity (Xue et al. 2015; Benazzo et al. 2017; Robinson et al. 2018; Grossen et al. 2020). Therefore, understanding the complex relationship between genetic diversity, inbreeding, genetic load, and demographic history are key to leveraging this information for management and conservation strategies of populations in the wild.

In large and outbred populations, strongly deleterious recessive alleles are “hidden” from selection in heterozygous form, making large populations more prone to experiencing inbreeding depression after going through a harsh bottleneck (Robinson et al. 2018; Kyriazis et al. 2021). On the other hand, small and isolated populations tend to have a higher burden of deleterious variants as a result of inbreeding (Robinson et al. 2018; Hedrick et al. 2019; Grossen et al. 2020; Khan et al. 2021; Stoffel et al. 2021). Accumulation of long ROH in the genome, indicators of inbreeding, is associated with increased presence of deleterious homozygotes and potential fitness effects (Szpiech et al. 2013; Szpiech et al. 2019; Stoffel et al. 2021; Steux and Szpiech 2023; Swinford et al. 2023).

In this work, we have generated five whole-genome sequences of K’gari dingoes to investigate the consequences of isolation and inbreeding on this island population. We find K’gari dingoes have lower genetic diversity and higher levels of runs of homozygosity compared to dingoes in the mainland and canids from large populations. Furthermore, we find a higher burden of deleterious homozygotes in K’gari dingoes compared to canids from large and stable populations, such as wolves and village dogs (fig. 4A).

Although the levels of ROH and estimated inbreeding found in K’gari dingoes (F_ROH_ = 0.59 on average; fig. 3) in some cases surpass those observed in wild populations of other species already experiencing inbreeding depression (supplementary table S3) (Kardos et al. 2018; Robinson et al. 2019; Khan et al. 2021; Stoffel et al. 2021; Taron et al. 2021; Colpitts et al. 2022), surveillance and necropsy records from the Queensland Parks and Wildlife Service (QPWS) and the latest report on the genetic health of K’gari dingoes (Allen 2023) have not recorded abnormal phenotypes in K’gari dingoes. Nonetheless, whether a small population will experience inbreeding depression or not depends on various factors, including the size of the ancestral population and the magnitude and history of bottleneck events (Kyriazis et al. 2021).

It has been observed that long-term small populations (i.e., populations that experienced early bottlenecks and have maintained small population size following the bottlenecks) can persist despite showing high levels of inbreeding by purging strongly recessive deleterious variants (Xue et al. 2015; Benazzo et al. 2017; Robinson et al. 2018; Grossen et al. 2020; Dussex et al. 2021), whereas populations that have gone through multiple recent bottlenecks will most likely experience inbreeding depression despite purging (Khan et al. 2021; Stoffel et al. 2021). At least two bottlenecks in the form of large-scale culling events have been documented during the past 120 years (Petrie 1995; Allen et al. 2015). Moreover, according to surveillance records from QPWS, dingoes in K’gari have maintained a small population size (<200 individuals) for the past 30 years, and evidence of inbreeding depression has not been observed. Our assessment of purging in this population suggests that while strongly deleterious alleles are likely being purged, there may be little to no purging of mildly deleterious alleles in this population. We hypothesize purging of strongly deleterious alleles may have played a role in the persistence of K’gari dingoes, as there is no current overt evidence of inbreeding depression despite the population having experienced recent severe bottlenecks. However, the long-term viability of this population is uncertain given its small population size and genetic isolation.

Current conservation and management programs treat dingoes as a single homogeneous population, although previous research has identified at least three distinct dingo populations (Savolainen et al. 2004; Ardalan et al. 2012; Oskarsson et al. 2012; Sacks et al. 2013; Cairns and Wilton 2016; Cairns et al. 2017; Greig et al. 2018; Zhang et al. 2020; Cairns et al. 2023). Adding to previous evidence, we find two distinct populations in the mainland (northwest and southeast), and one offshore in K’gari. Cairns et al. (Cairns et al. 2017) found dingoes in K’gari share the paternal haplogroup from northwest mainland dingoes but carry the maternal lineage from dingoes in the southeast. Consistent with these findings, we identify long IBD segments shared between dingoes from K’gari and northwest Australia, and between dingoes from K’gari and the southeast (supplementary fig. S2). In light of these findings, recognizing several dingo subpopulations may help inform conservation strategies.

This work is the first step in broadening our knowledge and understanding of the landscape of genetic diversity and deleterious variation in K’gari dingoes in order to better inform conservation and public policies. Our work, although limited by a small sample size and the sampling of several closely related samples, uses standard genomic analysis tools and indicates that the dingo population in K’gari is challenged by inbreeding as a result of founder effects and successive bottlenecks. We find the first evidence that K’gari dingoes may be purging strongly deleterious mutations, whilst accumulating a higher genetic load of mild/slightly deleterious mutations. Further research is needed to investigate the evolutionary history and genetic load of dingoes in K’gari using a larger sample size, which in turn, would allow us to incorporate deleterious variation to conduct genomic simulations to learn more about the future health of this population. In addition to this, dedicated monitoring for phenotypic or reproductive abnormalities such as reduced fertility, increased stillbirths, and congenital birth defects is recommended, as biological monitoring for symptoms of inbreeding depression is currently understudied in K’gari dingoes. QPWS and Butchulla Prescribed Body Corporates are left with the challenge of preserving dingoes in K’gari, balancing the need to address human safety concerns, and protecting the dingo population’s genetic health and viability into the future.

## MATERIALS AND METHODS

### Sampling and sequencing

In this study we analyze whole-genome sequencing data from a total of 41 canids, including 18 dingo samples from both K’gari (6) and mainland Australia (4 from the Southeast, and 8 from the Northeast). We also include 2 New Guinea singing dog samples, 11 gray wolf samples, 1 red wolf sample, and 9 village dog samples (supplementary table S2). We generated whole-genome data for five of the six K’gari dingo individuals analyzed in this study. Biological samples from these five individuals were collected between 2011 and 2013 from regions along the east coast of K’gari, including Cathedral Beach, Eurong, and Dundubara, and were sequenced using Illumina Hiseq 2000. Newly sequenced raw reads from K’gari dingoes and publicly available NCBI SRA raw reads were trimmed based on quality and adapter sequences using Trimmomatic v0.36 (Bolger et al. 2014) using the following parameters: ILLUMINACLIP:adapter.fa:2:30:10 SLIDINGWINDOW:4:20 MINLEN:5. Reads from all 41 canids were aligned to the CanFam3 reference genome using BWA-MEM v0.7.13 (Li and Durbin 2009) with parameters set as follows: -M -t 8 -r “ID:sampleID\tSM:sampleName”. Default parameters were used to index the reference genome prior to alignment. Mapped reads were converted from sequence alignment/map (SAM) format to binary alignment/map (BAM) format using the view command of samtools v1.3.1 (Danecek et al. 2021) and the following parameters: -@ 8 -b -F 4. The resulting BAM file was sorted based on chromosomes and alignment position with parameters set as follows: -@ 10 -o, and subsequently, indexed using default parameters. The sorted reads were used for marking PCR duplicates using the MarkDuplicates tool of the Picard toolset v2.25.2 (Broad Institute 2019) with default parameters.

### Genotype calling, filtering, and phasing

All samples were jointly genotyped with the Genome Analysis Toolkit (GATK) v4.2 (McKenna et al. 2010). GATK HaplotypeCaller was used to call genomic variants with parameters set as follows: -Xmx5g -Xms5g -XX:ParallelGCThreads=4 -ERC GVCF. Samples were merged using GATK CombineGVCFs with the following parameters: - Xmx5g -Xms5g -XX:ParallelGCThreads=4. Lastly, GATK GenotypeGVCFs was used with default parameters to call the variants.

The resulting VCF file containing raw genotype calls for all 41 canids was then filtered using BCFtools v1.18 (Danecek et al. 2021) in the following way: Only sites with biallelic snps and only sites with a site-level quality score (QUAL > 50) were retained. Any individual genotype with (GQ <= 19) was set to missing. All sites with >= 10% missing data were removed. All sites with no variation among the 41 samples (i.e., monomorphic sites) were removed. This resulted in 3,745,051 sites. Samples were then split into three separate VCF files: all wolves (n=12), all village dogs (n=9), and all dingoes plus NGSD (n=20). Each group-level VCF was filtered separately, in the following way. As a result of small sample sizes, formal statistical tests for excess heterozygosity have no power. Therefore, to filter for excess heterozygosity, only sites where heterozygote calls made up < 75% within each species-level VCF were retained. This resulted in 3,729,028 sites for wolves, 3,649,859 sites for village dogs, and 3,733,743 sites for dingoes plus NGSD. Finally, all samples were merged back together by intersecting all sites among the three data sets, resulting in 3,632,373 sites.

SHAPEIT2.v904 (Delaneau et al. 2014) was used with default parameters to phase the genotypes of all 41 canids, using the pedigree-based recombination map from Campbell, et al. (Campbell et al. 2016). The ancestral and derived states of each variant for the 18 dingo samples was taken from Kumar et al. (Kumar et al. 2023).

### Relatedness and Population Structure

We used BCFtools v1.18 (Danecek et al. 2021) to remove sites with 2 or fewer minor allele counts in the 18 dingo samples, which resulted in 887,759 biallelic loci. We then used ASD v1.1.0a (Szpiech 2014) with the flag --ibs to compute the proportion of sites at which a pair of individuals shared 0, 1, and 2 alleles Identical-By-State (IBS) for each pair of dingoes.

To assess relatedness among K’gari dingoes we plotted IBS0 (the proportion of 0 identical alleles at a given position being shared by two individuals) vs IBS2 (the proportion of 2 identical alleles being shared by two individuals). Based on records from Queensland Parks and Wildlife Service (QPWS), there are three samples in the K’gari dingo population that look to be related to one another by first-degree relationship (i.e., samples Y2, B44, and Y4). To verify these relationships, we used KING v2.3.1 (Manichaikul et al. 2010) with the --kinship flag, which incorporates an algorithm that estimates a kinship coefficient between pairs of individuals independent of the population’s allele frequencies.

To assess population structure among canids we computed pairwise allele sharing dissimilarities (Szpiech 2014) for all 41 canids included in this study and decomposed the dissimilarity matrix with multidimensional scaling (MDS). For MDS analysis using only the 18 dingo samples, we subset the allele sharing dissimilarities from the 41 canids and recomputed the MDS.

### Genetic diversity within and between canid populations

We estimated allelic richness, private allelic richness, and the number of alleles private to pairs of populations (found in each population but no other) using ADZE v1.0 (Szpiech et al. 2008). ADZE uses a rarefaction approach to compare these statistics at standardized sample sizes. For these analyses we grouped the two mainland dingo populations (northwest and southeast) and excluded the red wolf and NGSD samples because of small sample sizes. Parameters were set as follows: COMB 1, K_RANGE 2, and MAX_G 24. To visualize results, we used R (R Core Team 2022) to plot the mean of each estimate calculated for each standardized sample size.

### Estimating IBD and ROH

To call IBD tracts, we used RefinedIBD v17Jan20.102 (Browning and Browning 2013) to identify Identical-By-Descent (IBD) segments with default parameters and the CanFam3.1 genomic map (Campbell et al. 2016). We called runs of homozygosity (ROH) in each canid population using GARLIC v1.6.0a (Szpiech et al. 2017). Parameters were set as follows: --winsize 100, --auto-winsize, --auto-overlap-frac, -- size-bounds 1000000 3000000 5000000, --gl-type GQ, --resample 40. This results in 4 length classes of ROH: <1Mbp, ≥1Mbp to <3Mbp, ≥3Mbp to <5Mbp, and ≥5Mbp.

As GARLIC uses allele frequency information to call ROH, we account for the close relatedness among the K’gari dingo samples, which could result in miscalling ROH in these individuals, by using the best linear unbiased estimator for allele frequencies (McPeek et al. 2004) computed from pooling all dingoes together. The allele frequency at locus *i*, *f_i_*, is computed as 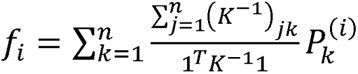, where 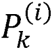 is the fraction of derived alleles at a locus in individual *k*, *K* is a symmetric matrix of kinship coefficients, *K*^-1^ is the inverse matrix of *K*, 1 is a column vector of *n* elements with all entries equal to 1, and 1*^T^* is the transpose of 1. The kinship matrix for dingoes obtained from the -- make-king option in PLINK (Purcell et al. 2007), and any negative entries were set to 0 (https://github.com/szpiech/blue_af). The allele frequencies were passed to GARLIC with the --freq-file option for use in calling ROH for the 18 dingoes and the 2 New Guinea singing dog samples.

Results were filtered to keep only the ROH segments longer than 0.5Mbp. We summarized the number of ROH segments (nROH) and sum total length of ROH (sROH) within each individual and each length class. The fraction of the genome within ROH was calculated as the total length of ROH of 0.5 Mbp or longer within an individual and divided by the total length of the CanFam3.1 autosomal reference genome (2,203,764,842 nucleotides). Similarly, we estimated inbreeding coefficients as given by ROH for each population (F_ROH_). This coefficient represents the proportion of the autosomal genome covered by ROH 1Mbp or longer.

### Assessing burden of deleterious variants

Variants called across all 41 canids were annotated with Ensembl Variant Effect Predictor (VEP) from release 83 (McLaren et al. 2010). We used SIFT 4G (Vaser et al. 2016) to predict whether an alternate allele at a coding mutation is tolerated or deleterious for protein function based on the CanFam3.1 genome annotations. We sort these mutations into three categories: Deleterious, Tolerated, and Loss-of-Function (LoF). The “Deleterious” group includes only nonsynonymous variants that were predicted as damaging (1,921 sites). The “Tolerated” group contains all synonymous and nonsynonymous variants predicted as tolerated (16,380 sites). If a mutation has more than one prediction we retain the most damaging prediction, i.e., if there is at least one damaging prediction among multiple predictions, it is classified as “Deleterious”. The LoF category is composed of mutations that are annotated as “Stop-Gain”, “Stop-Loss”, or “Start-Loss” (146 sites) and is a separate category from those predicted as “Deleterious”.

We generated a second set of SIFT predictions, this time, requiring that the derived allele is the one with the prediction. For comparison purposes, we kept both mainland dingo populations (northwest and southeast) together. We created the same three categories (1,357 Deleterious, 7,713 Tolerated, and 88 LoF sites) and counted the number of derived-allele genotypes per individual for each category by summing the number of heterozygotes plus two times the number of homozygotes.

To assess purging of deleterious variants we compute the Rxy statistic (Xue et al. 2015), which compares the derived allele frequencies of one set of variant sites (e.g., putatively deleterious sites) normalized by a set of putatively neutral sites (e.g., intergenic sites) between two populations X and Y. We set X = K’gari dingoes and Y = Mainland dingoes, and considered several sets of sites, including SIFT-classified Deleterious, SIFT-classified Tolerated, LoF, and sites annotated as synonymous (n = 4708), nonsynonymous (n = 4668), or intergenic sites (n = 100,000). For each of these sets of sites, we normalize against a second (independent) set of intergenic sites (n= 100,000). To account for the close relatedness among K’gari dingoes we implement a kinship-weighted Rxy statistic. The modified Rxy statistic (https://github.com/szpiech/Rxy-kin) incorporates the kinship matrix for K’gari and mainland dingoes obtained from the --make-king option in PLINK (Purcell et al. 2007), where any negative entries are set to 0. The statistic is defined (Xue et al. 2015) as 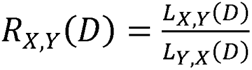, where 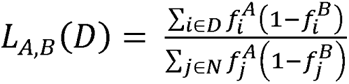, D is a set of test sites, N is a set of putatively neutral sites, 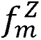 is the derived allele frequency at site m in population Z. To account for relatedness when estimating allele frequencies, *f_i_* are computed as described above, using weights derived from the kinship matrix. We estimate the variance in our kinship-weighted Rxy statistic using a jackknife sampling of 10,000 iterations with 20% reduction per iteration.

## DATA AVAILABILITY

Raw sequence data of each of the K’gari dingo genomes generated in this study are available on SRA under project PRJNA1021344.

## ACKNOWLEDGEMENTS

The authors would like to acknowledge the Butchulla People as the traditional custodians of K’gari. Thanks to the Queensland Parks and Wildlife Service for access to the K’gari wongari (dingo) DNA sample collection, and thanks to Anna María Calderón for helpful feedback on early versions of this work. This work was supported by the National Institute of General Medical Sciences of the National Institutes of Health under Award Number R35GM146926 (AVLA and ZAS), by start-up funds from the Pennsylvania State University’s Department of Biology (AVLA and ZAS), and by a grant from the Australian Dingo Foundation (KMC). Computations for this research were performed using the Pennsylvania State University’s Institute for Computational Data Sciences’ Roar supercomputer.

## SUPPLEMENTARY MATERIAL

Supplementary data are available at XXXX.

