## Supplementary figures and images for "Genomic consequences of isolation and inbreeding in an island dingo population"

### supplementary fig. S1

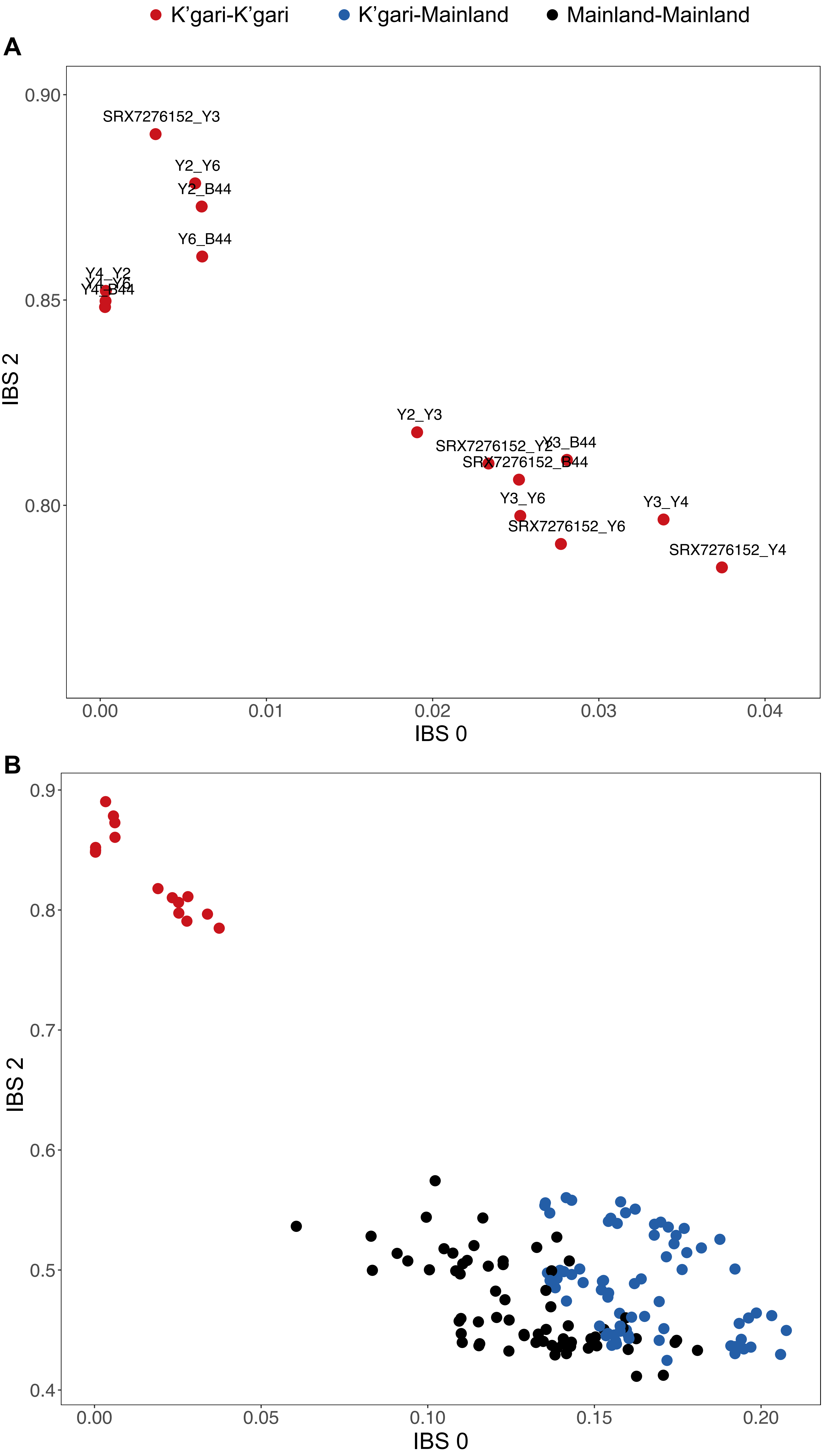

### supplementary fig. S2

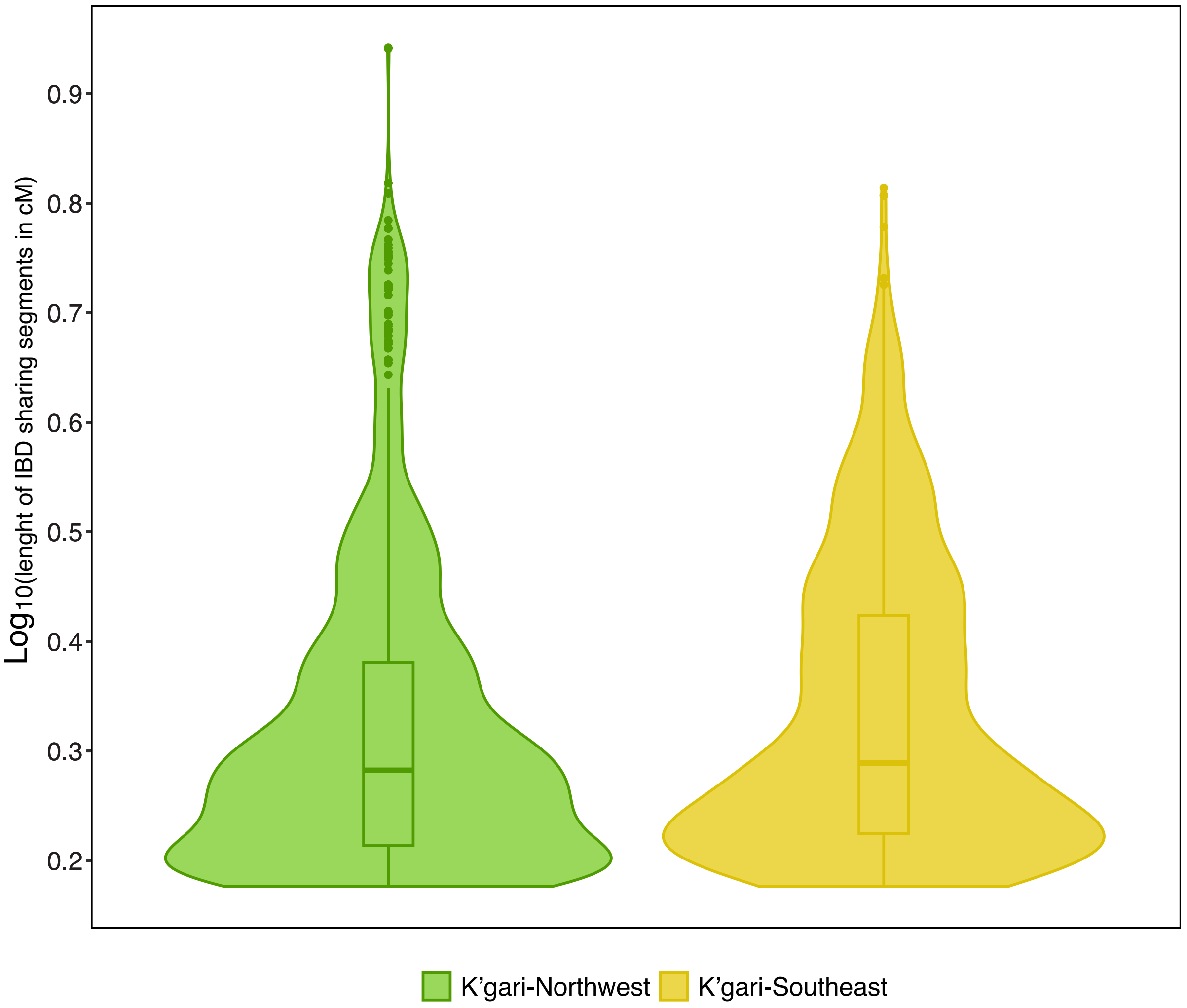

### supplementary fig. S3

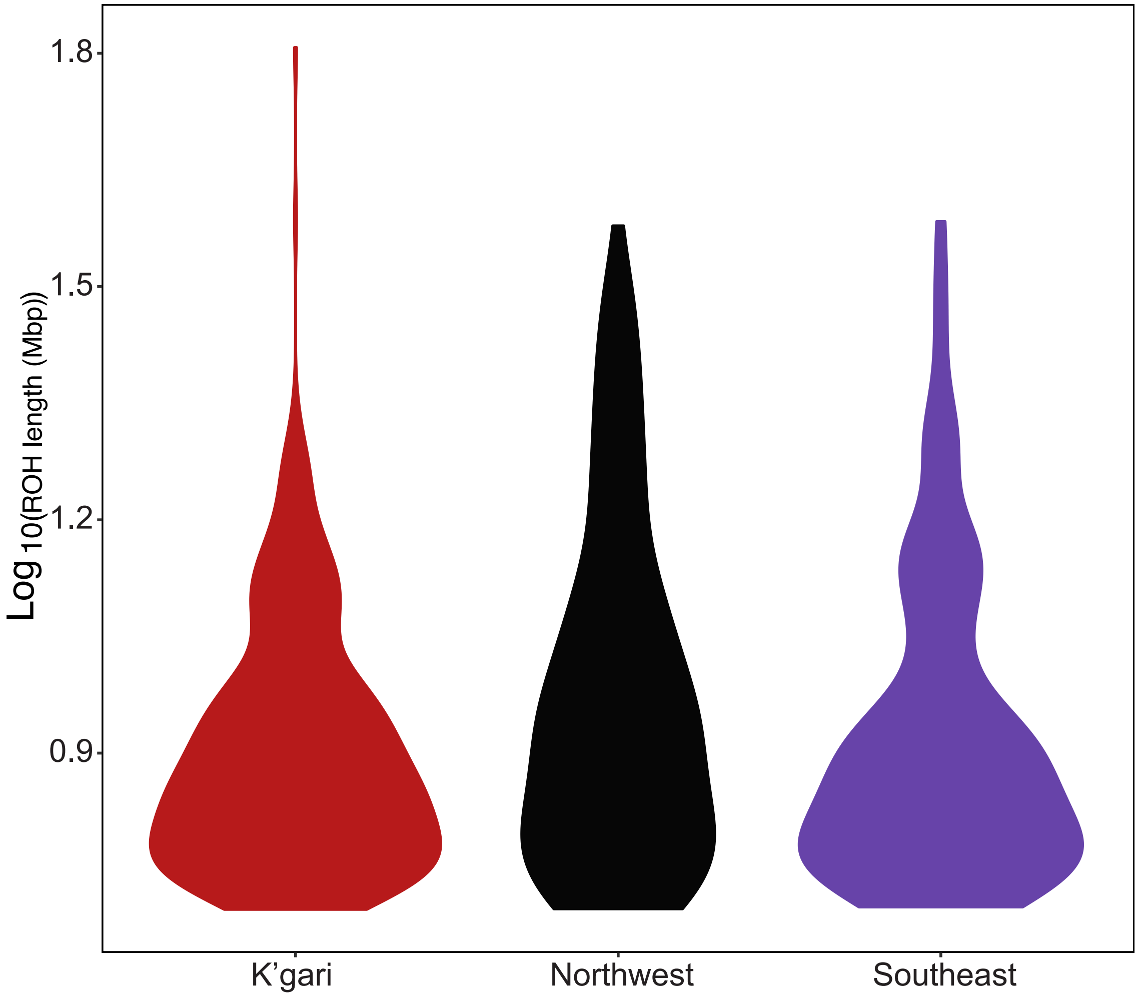

### supplementary fig. S4

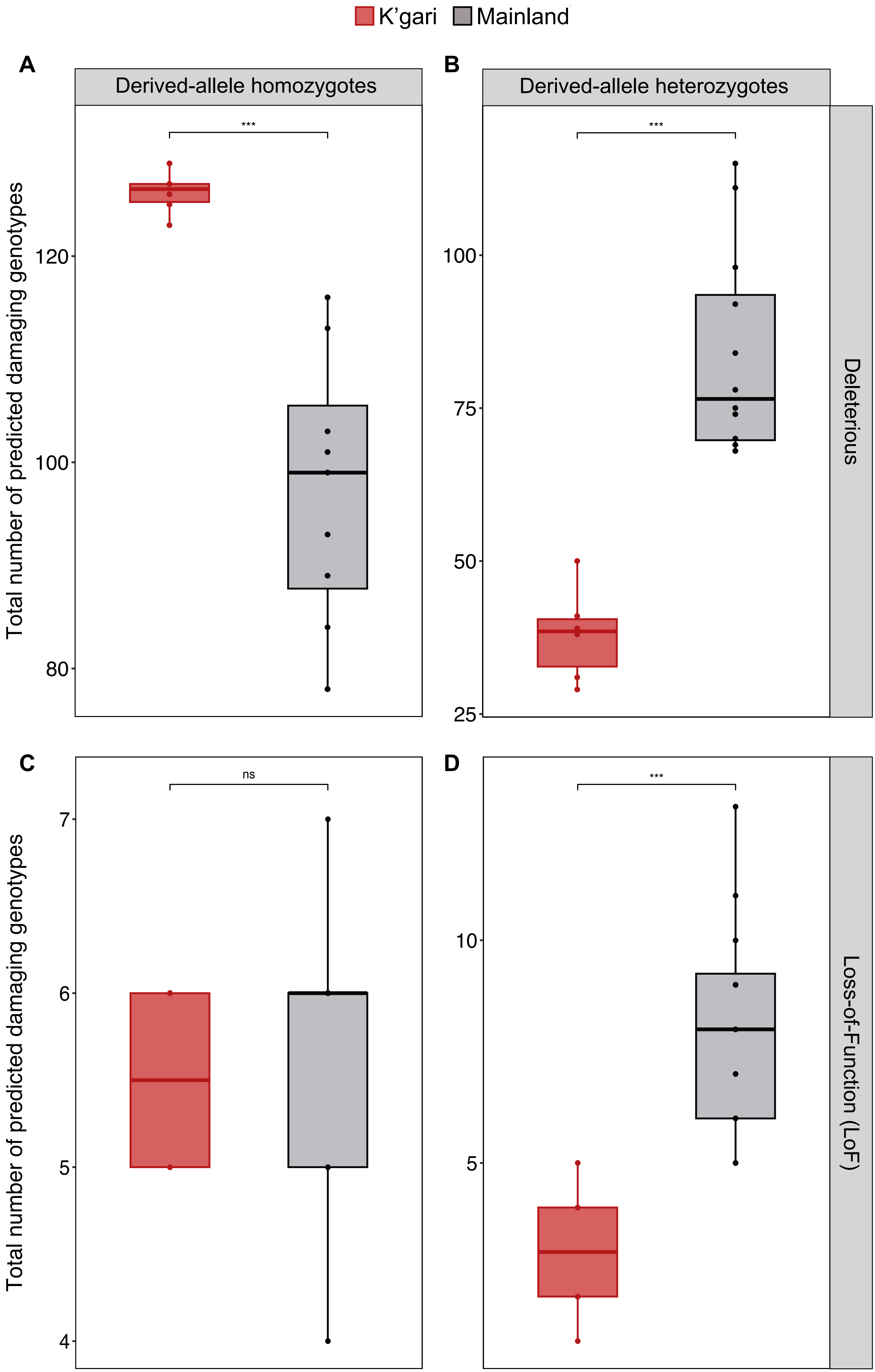

### supplementary fig. S5

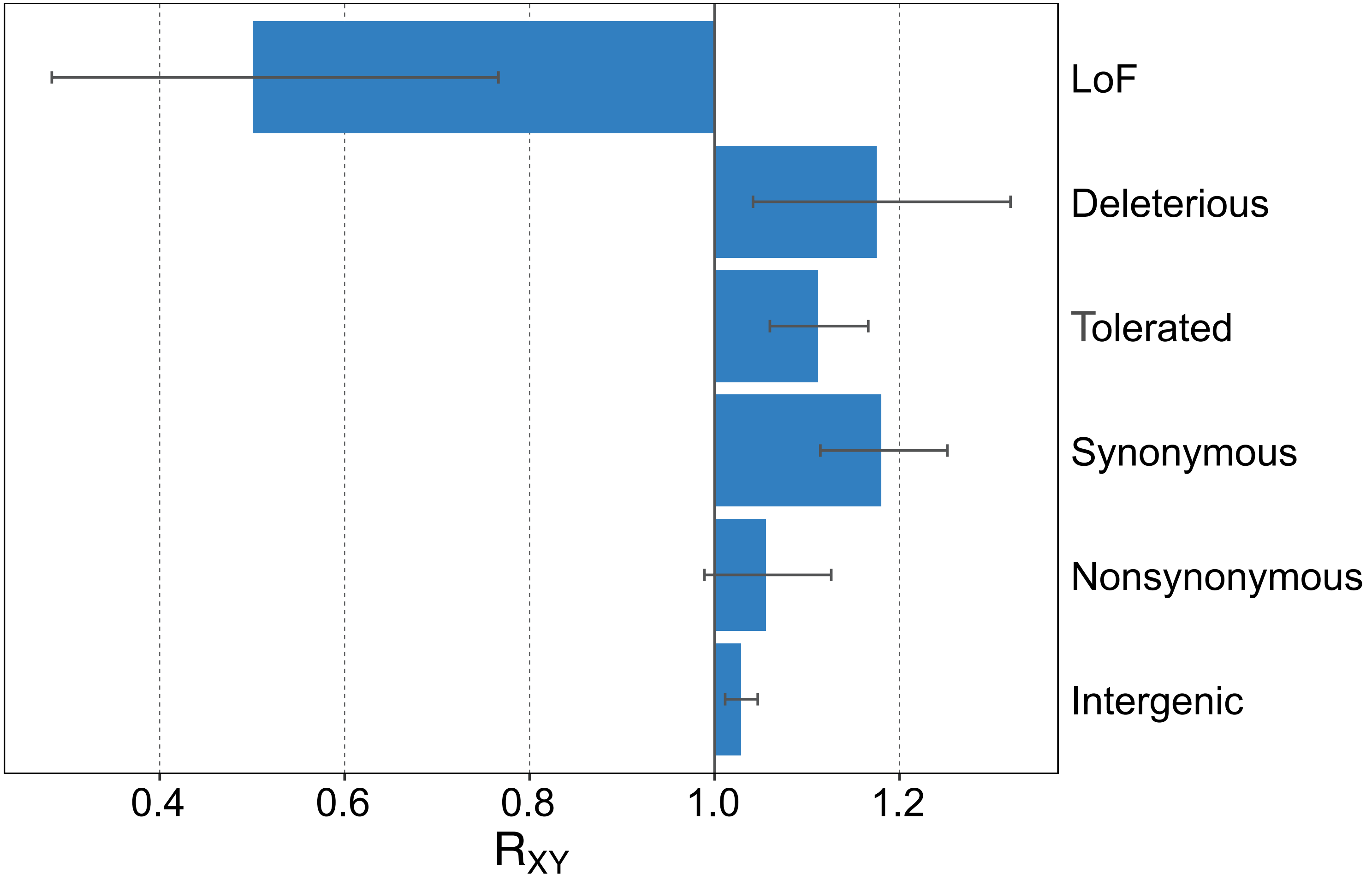
